# CO-EXPRESSION OF DISTINCT COILED COILS CAN LEAD TO THEIR ENTANGLEMENT

**DOI:** 10.1101/2023.05.17.541026

**Authors:** Nikola A. Mizgier, Charlie E. Jones, Anthony V. Furano

## Abstract

L1 (LINE1) non-LTR retrotransposons are ubiquitous genomic parasites and the dominant transposable element in humans having generated about 40% of their genomic DNA during their ∼100 million years (Myr) of activity in primates. L1 replicates in early embryos, causing genetic diversity and defects, but can be active in some somatic stem cells, tumors and during aging. L1 encodes two proteins essential for retrotransposition: ORF2, a replicase, and ORF1, a coiled coil mediated homo trimer, which functions as a nucleic acid chaperone. Both proteins contain highly conserved domains and preferentially bind their encoding transcript to form an L1 ribonucleoprotein (RNP), which mediates retrotransposition. However, the coiled coil has undergone episodes of substantial amino acid replacement to the extent that a given L1 family can concurrently express multiple ORF1s that differ in the sequence of their coiled coils. Here we show that such distinct ORF1p sequences can become entangled forming heterotrimers when co-expressed from separate vectors.

## INTRODUCTION

L1 non-LTR retrotransposons, which replicate by copying their RNA transcript into genomic DNA, reside in most eukaryotic genomes (1) and are active in nearly all vertebrates where they share the same general structure: a 5’ untranslated region (UTR), which has regulatory functions; two protein encoding sequences (ORF1 and ORF2) and a 3’ UTR (Figure 1, top panel) (2). All vertebrate ORF1 proteins (ORF1p) contain a coiled-coil domain shown in mouse and humans to mediate trimerization of the ORF1p monomer, which is required for high affinity binding to single stranded nucleic acid (ssNA) and NA-chaperone activity that are essential for retrotransposition (3-6). ORF2 encodes the replicase, which contains highly conserved reverse transcriptase and endonuclease domains (7-9).

**Figure 1.**
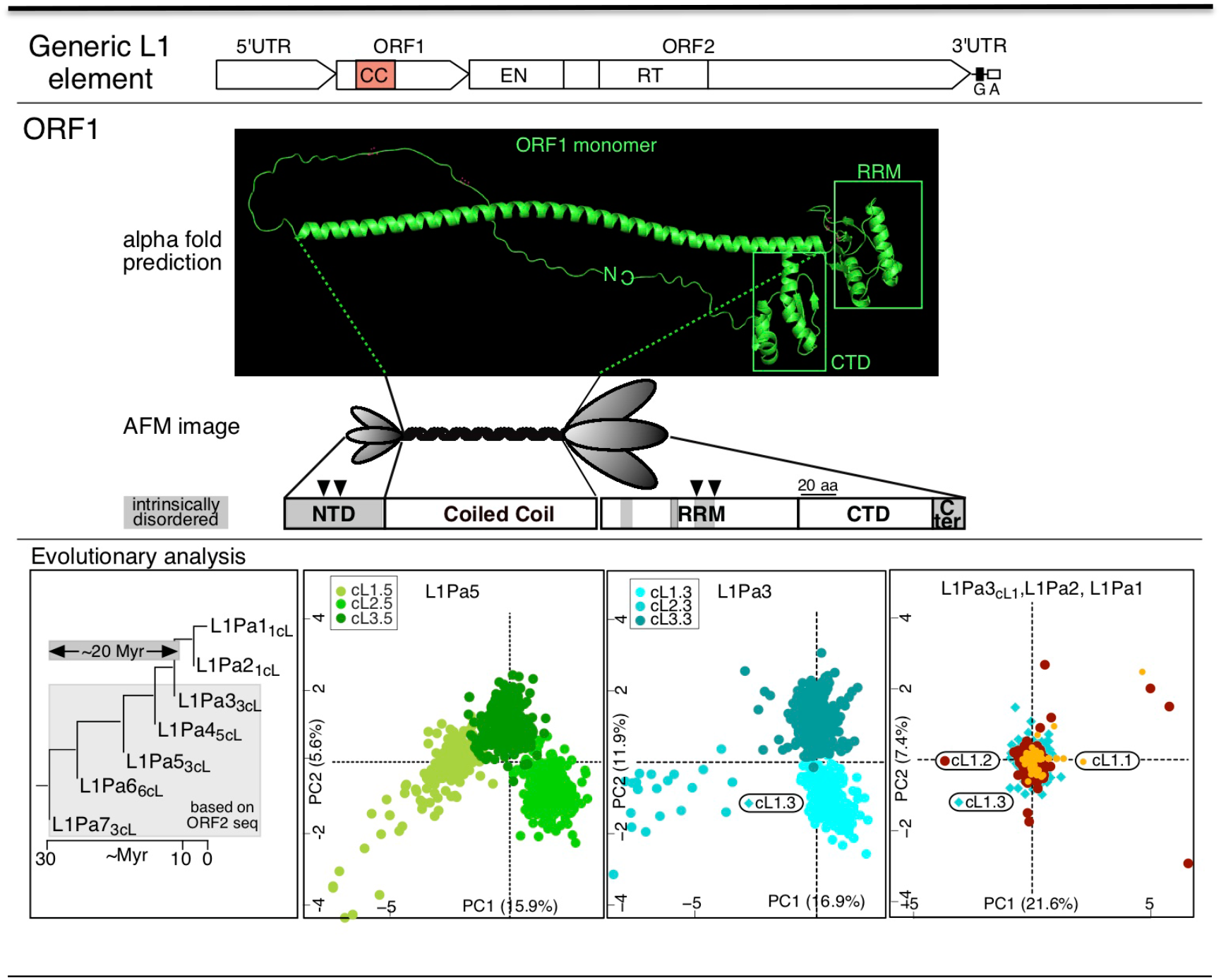
ORF1 structure and coiled coil variants. The top panel shows a generic L1 element depicting to scale the coiled coil domain (CC) in ORF1, the endonuclease (EN) and reverse transcriptase (RT) domains in ORF2, and the G-rich (G) and A-rich (A) motifs in the 3’ UTR (2, 13). The middle ORF1 panel shows the AlphaFold predicted structure for the ORF1p monomer, https://alphafold.ebi.ac.uk/entry/Q9UN81 (14), and below that a rendering of the atomic force microscopic (AFM) image of the ORF1p trimer (15), its predicted secondary structure (α helix, β sheet, white boxes) and intrinsically disordered regions, grey boxes (16), and essential phosphorylation sites (arrow heads) (17) - adapted from Figure 1A of (18). The bottom Evolutionary analysis panel was adapted from Figure 3 of (13). The number of distinct coiled coils (which present as clusters of related divergent sequences, ncL – see text) is indicated for each family on the phylogenetic tree that was derived from ORF2 sequences shown in the left most panel. Cluster 3 of L1Pa5 (cL3.5) encodes 555p.

Continued emergence, amplification and senescence of successive L1 families accompanied the evolution of most mammals. Each family generated hundreds to many thousands of copies that were retained in the genome. Most were defective upon insertion or not being under purifying selection have decayed by the accumulation of random mutations. Although the sequences of these copies have diverged from their once active progenitor, its sequence can be inferred from these fossils, which thereby provide a record of the evolutionary antecedents of the modern families active in present day species, which in humans is designated L1Pa1 (or L1Hs). L1Pa1 is the product of a primate-specific L1 lineage that emerged 80-120 million years ago (MYA) and consists of a largely single lineage of ∼16 distinct L1 clades (L1Pa16 – L1Pa1) based on ORF2 and 3’ UTR sequences (10, 11).

In contrast to most of the L1 sequence, the 5’ UTR and amino-terminal half of ORF1 are evolutionarily labile (2, 12), especially the coiled coil that has been subject to repeated episodes of extensive amino acid substitution(13). The last such episode in primates lasted for ∼20 Myr during the evolution of L1Pa7-L1Pa3 (*i*.*e*., from ∼30–10 MYA, Figure 1, bottom left panel). Unexpectedly, each L1 family that was active during this episode concurrently expressed three or more distinct ORF1ps based on their coiled coil sequences, which due to sequence divergence, are now represented by clusters (cL) of related sequence that can be resolved by metric multidimensional scaling (MMDS) (13).

The left most bottom panel of Figure 1 shows the number of coiled coil clusters (ncL) in the L1Pa7–L1Pa1 families detected by MMDS, and the others show the results of cluster analyses for the L1Pa5, L1Pa3, L1Pa2 and L1Pa1 families. The latter two families diverged about 7 MYA and the consensus coiled coil sequences derived from each cluster are identical and marked the end of this episode of coiled coil change. When active, the L1Pa5 family expressed 3 ORF1 sequences arbitrarily designated: cL1.5, cL2.5 and cL3.5. Consistent with work by others (19-24) we proposed and provided experimental evidence indicating that the enlarged functional coiled coil sequence space can buffer the effect of inactivating epistatic mutations in the coiled coil (13).

These findings also raised the question of whether co-expressed distinct coiled coils, which could have occurred when L1Pa5 was replicatively active, could become entangled and form heterotrimers. A recent proteome-wide ribosome profiling study showed that coiled coil formation between nascent peptides on neighboring (*i*.*e*., cis) ribosomes was the most common mechanism that mediated homomer (dimer) formation of numerous proteins(25). Also, they found no evidence for *trans*-mediated interactions (*i*.*e*., between nascent peptides undergoing translation on separate mRNAs) either on a proteome level search or when examined experimentally by testing for interaction between the lamin C 350 amino acid “rod” region, 320 amino acids of which form 3 coiled coils connected by short linker sequences (26). In the case of distinct L1Pa5 coiled coil sequences, entanglement would have to not only be mediated in *trans*, but also involve α-helices that differed in primary sequence.

## Results and Discussion

To provide the simplest experimental test for entanglement we fused either Flag (FG) or HA (a peptide derived from the human influenza antigen) epitopes to the C-terminus of various ORF1 coding sequences and expressed them in HEK293F cells (27) using a pcDNA3-based expression vector as described in the **Materials and Methods** and the legends to the various figures. To prevent non-specific aggregation of trimers in cell lysates all buffers contained 10 μM of a 52 nt deoxynucleotide of T (dT_52_)(3), see Materials and Methods.

Panel A of Figure 2 shows the amino acid sequences of the NTD and coiled coil of 111p, 151p and 555p, which are encoded respectively by the modern L1Pa1-ORF1, a mosaic L1Pa1/L1Pa5-ORF1, and L1Pa5-ORF1 resuscitated from the now extinct ancestral L1Pa5 family (corresponds to cL3.5, Figure 1). The purified proteins are essentially indistinguishable with respect to ssNA affinity, and a FRET based NA chaperone assay (annealing/strand exchange) (3, 28, 29). However, whereas 555p displays about 80% of the 111p activity in a cell culture based retrotransposition assay the mosaic 151p is inactive due to its inability to form tightly compacted complexes on ssNA (29).

**Figure 2.**
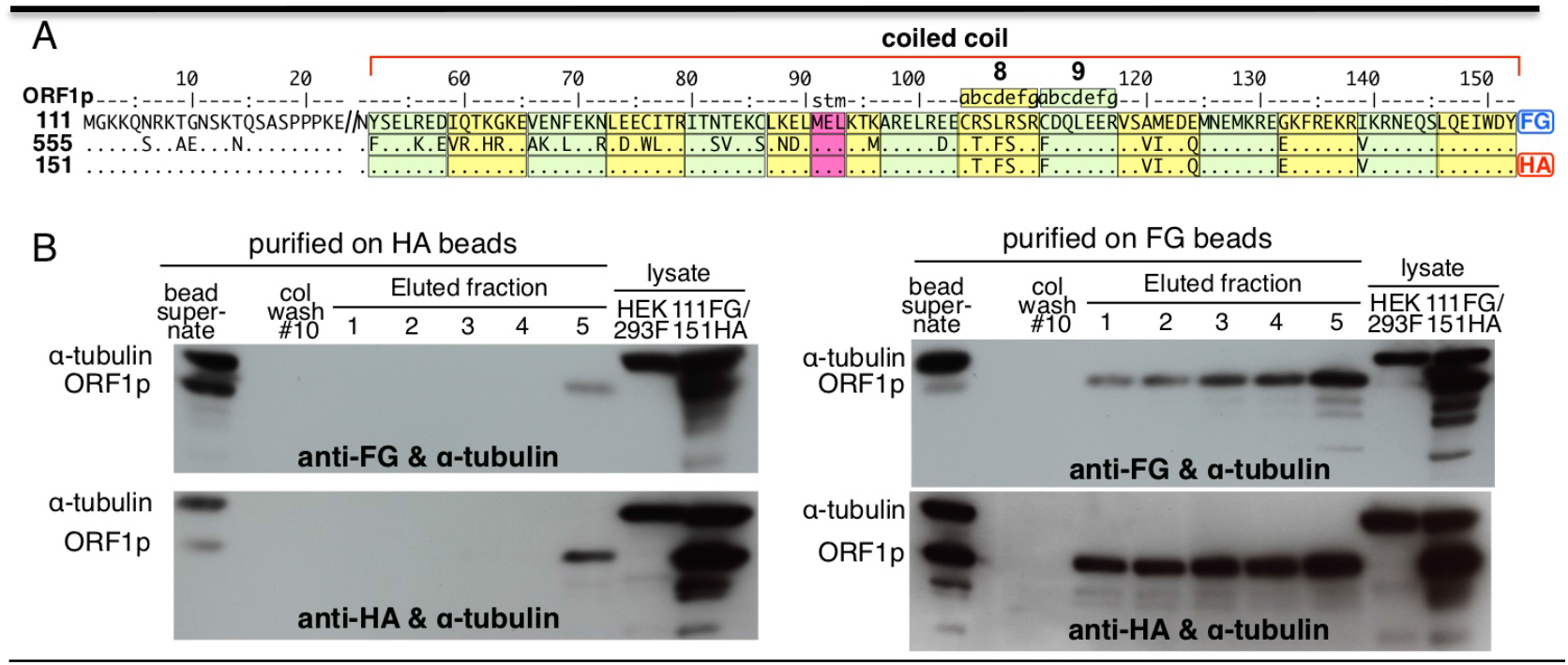
Entanglement of coiled coils that differ at nine positions. A. Alignment of 555p and 151p to the 111p reference sequence. FG and HA refer respectively to the Flag and HA peptide epitope tags, dots indicate identity. B. Western blots of proteins that were captured by anti-HA (left) or anti-FG agarose beads (right) from cleared lysates prepared from HEK293F cells that had been co-transfected with pcDNA3.1(+)-puro based vectors expressing either ORF1p-111-3XFG (111-FG) or ORF1p-151-HA (151-HA) as described in the Materials and Methods. The upper blots were challenged with anti-FG and the lower blots with anti-FA and both with anti α-tubulin. In each instance lanes 1-10 of the blotted gel, starting at the left were loaded with: (1) the bead supernatant, (2) stained protein marker, (3) column wash #10, (4-8) eluted fractions 1-5, and lanes 9 & 10 respectively, cleared lysate from non-transfected HEK293F and the doubly transfected cells. See Material & Methods, *Protein purification on affinity agarose beads* for the meaning of these sample designations.

The 111-FG and 151-HA proteins differ at 9 coiled coil positions (Figure 2A) and cleared lysates from HEK293F cells that had been co-transfected with expression vectors for these proteins were incubated with anti-FG or anti-HA agarose beads (hereafter beads). After elution of bound proteins by their cognate epitopes, the eluates were denatured and subject to electrophoresis, under denaturing conditions. This treatment converts the ∼120 kD ORF1p timers to their constituent ∼40 kD monomers. The gels were blotted to filters and exposed to anti-FG or anti-HA antibodies as described in the **Material and Methods**. Figure 2B shows that protein recovered from either the HA-beads (left panels) or FG-beads (right panels) contained both ORF1p-FG monomers and ORF1p-HA monomers (top and bottom panels respectively).

Figure 3 shows the results when co-expressed proteins differ by 21 coiled coil positions were purified on just anti-FG beads, which bind its epitope far more efficiently than the anti-HA beads (*cf*. left and right panels in Figure 2). Figure 3B shows that both HA-tagged and FG-tagged ORF1p monomers were recovered from the anti-FG beads.

**Fig 3.**
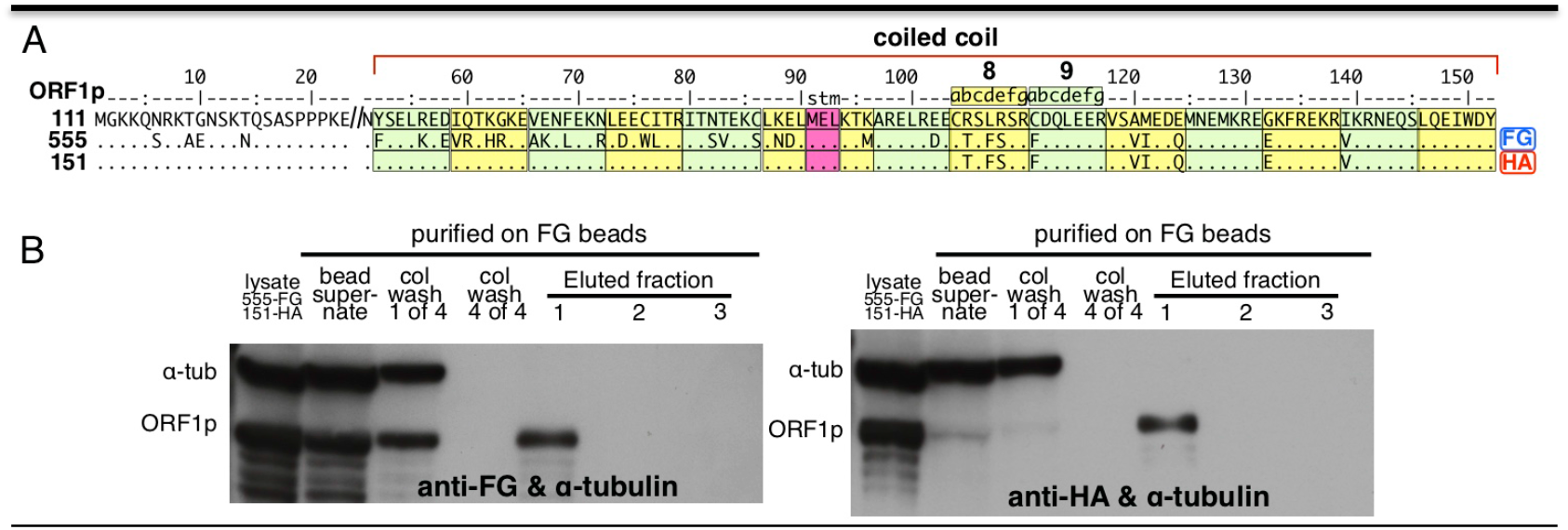
Entanglement of coiled coils that differ at twenty-one positions. A. Alignment of 555-FG and 151-HA to 111p, showing the twenty-one differences between 555p and 151p. B. Western blots of proteins captured by anti-FG agarose beads from cleared lysates prepared from HEK293F cells that had been transfected with expression vectors for both 555-FG and 151-HA as described in the Materials and Methods. In this case the FG-bound proteins were recovered from the anti-FG beads by batch elution. The blots were challenged with either anti-FG or anti-HA, and in both cases by α-tubulin. In both instances lanes 1-7 of the blotted gel starting at the left were loaded with: (1) cleared lysate of the doubly transfected cells, lane (2) bead supernatant, lanes (3 & 4) respectively with column washes #1 and #4, lanes (5-7), eluted fractions 1-3. See Material & Methods, *Protein purification on affinity agarose beads* for the meaning of these sample designations.

Supplemental Figure 1 presents the results of control experiments that support the above conclusions and Figure 4 shows that co-expression is required for entanglement. As described in **Materials and Methods** cleared lysates of HEK293F that had been transfected with 111-FG or 151-HA were mixed for 2 hours at 25º and then challenged with anti-FG beads, the eluate of which was processed as described for the experiments in Figures 2 & 3. In contrast to the co-transfection results, only 111-FG was recovered from the anti-FG resin (*cf*., left and right panels of Figure 4), indicating that heterotrimer formation depends on co-expression.

**Figure 4.**
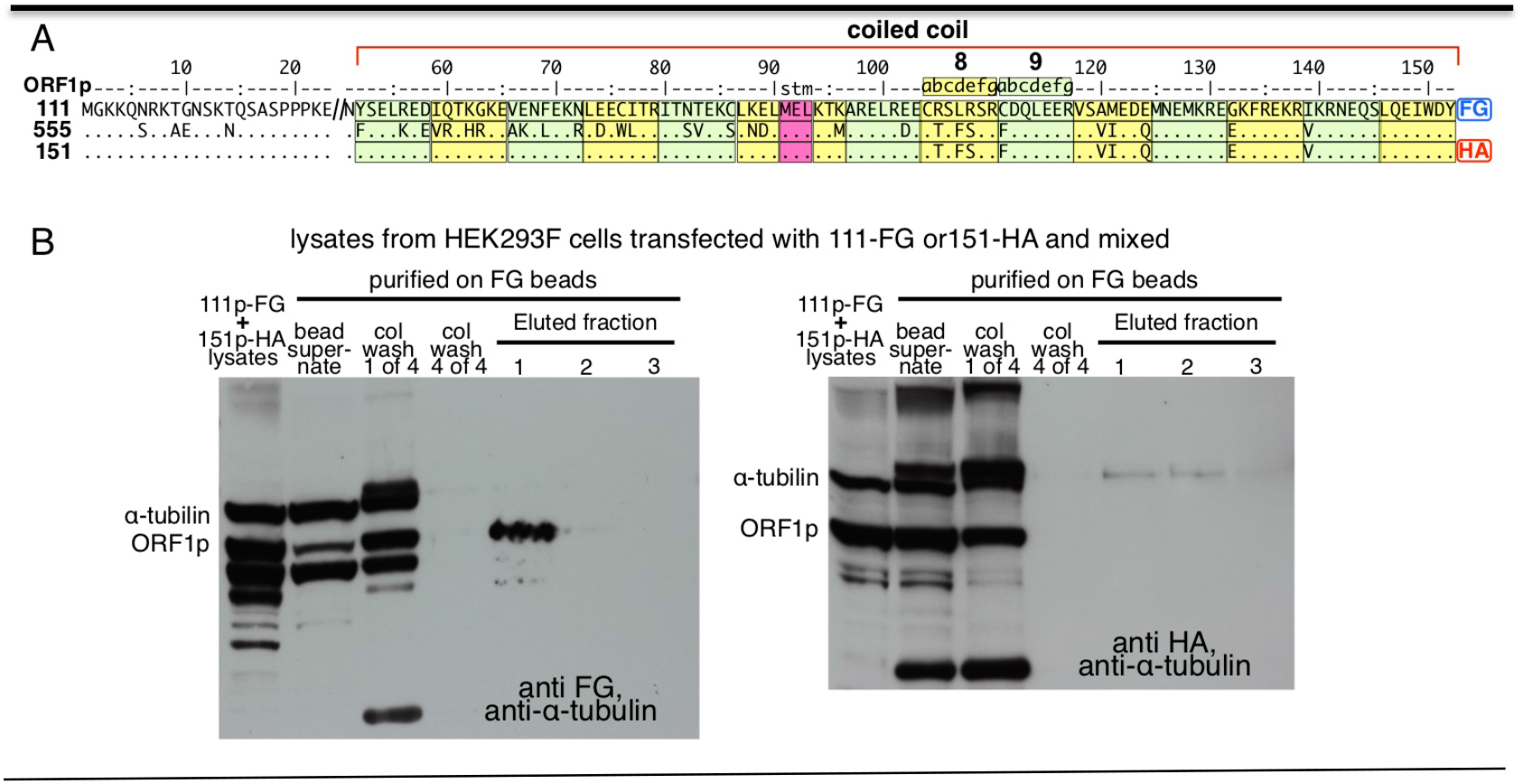
Entanglement requires co-expression. A. Alignment of 111p-FG and 151p-HA as described for Figure 2. B. Western blots of proteins captured by anti-FG agarose beads from cleared lysates prepared from HEK293F cells that had been transfected with expression vectors for either 555-FG or 151-HA and then mixed before processing as described in the *Mixed lysate experiment* section of the Materials and Methods (the lane designations are the same as for Figure 3). Again, the blots were challenged with either anti-FG or anti-HA, and in both cases by α-tubulin. In contrast to the co-transfection results, no HA-tagged proteins were recovered from the FG-beads.

Although these studies were prompted by our evolutionary evidence for concurrent expression of L1 elements encoding distinct coiled coils they do not speak to the issue of whether ORF1p expressed from full length L1 elements can also become entangled, and we have not attempted to pursue this experimentally. However, our findings indicate that the FG- and HA-tagged ORF1p are translated in close enough approximation to assemble hybrid trimers. Theoretical considerations (30) and experimental evidence (31-33) indicate that L1 proteins (ORF1p and ORF2p) bind to their encoding transcript, known as *cis* preference, to form an L1RNP retrotransposition intermediate. Here the meaning of *cis* is more general than and distinct from its use to describe the interaction of nascent peptides undergoing translation on ribosomes bound to the same transcript that leads to coiled coil mediated homomer (dimer) formation (25) or in the case of ORF1p, trimer formation.

As hundreds of trimers are present in a single L1RNP (15), it is difficult to visualize how trimers once formed by interaction of nascent ORF1p monomers emerging from neighboring ribosomes would then bind to their encoding transcript, as this would presumably block further translation. Additionally, Figure 2 showed that trimers can be assembled from monomers expressed from distinct ORF1 transcripts. Achieving this during co-translation would require interaction between nascent monomers emerging from ribosomes in *trans* (*i*.*e*., on distinct mRNAs). This phenomenon was not observed in a recent proteome wide study *in vivo* (25). However, there is evidence for transcript clustering, and proteins that comprise multi-protein complexes are often translated in intimate association referred to as “translation factories”, *e*.*g*., (34-36). In addition, undefined cytoplasmic ORF1p foci can be generated in mouse cells from endogenous L1 elements (37) and in human cells from ORF1p expression vectors and L1 retrotransposition reporter plasmids (38-40). In the latter case the foci include stress granules and P-bodies. Whether such foci reflect sites of protein synthesis that could account for co-assembly of ORF1p in *trans* is not known.

## Materials and Methods

### Plasmids and protein expression

We expressed various epitope-tagged versions of three ORF1 proteins, denoted 111p, 555p and 151p into the BamH1 and EcoR1 sites pORF1-Flag. As previously described (Supplementary Information, (17)), the expression vector was derived from pcDNA3.1(+)-Puro (from the Don Ganem laboratory, University of California San Francisco). In this construct the gene for neomycin resistance had been replaced by one for puromycin resistance. Because its DNA sequence had not been reported, we determined the sequence of the relevant region of the plasmid and deposited it to Addgene as pORF1-Flag. We had earlier added the FLAG® (Millipore Sigma) peptide (FG, DYKDDDDK) to 111p and 151p using the methods presented in the Supplementary Information of reference (17). Here we converted 111pFG to 111p3XFG or to 111pHA (YPYDVPDYA) and converted 151pFG to 151pHA using the NEBaseChanger tool. All DNA edits were verified by DNA sequencing. The ORF1p coding sequences are shown in Figure 2 of Supplementary Information (SI-CO-EXPRESSION OF DISTINCT COILED COILS CAN LEAD TO THEIR ENTANGLEMENT.pdf).

### Buffers and other reagents

TBS is 50 mM Tris, 150 mM NaCl. Where indicated this buffer was supplemented with 10 μM dT_52_ (a 52 nt deoxynucleotide of T), which eliminates aggregation of ORF1p trimers that can occur to varying extents at <0.5M NaCl (3). PBS is phosphate buffered saline (0.15 M NaCl). Cell lysis buffer is M-Per (ThermoFisher) adjusted to 0.15M NaCl, 10μM dT_53_, 5μg/ml leupeptin (Millipore Sigma), and 1 mini tablet of Pierce™ Protease Inhibitor (ThermoFisher) / 10 ml of buffer. Anti-FG M2 and anti-HA agarose beads were purchased from Millipore Sigma. 3XFG and HA peptides were purchased from Sigma-Aldrich and dissolved respectively at 100 μg/ml and 500 μg/ml in TBS.

### Protein expression

Proteins were synthesized in HEK293F (27) cells grown without serum in suspension using the medium and transfection reagents supplied in the GIBCO Expi293™ Expression System as described in its accompanying protocols. Generally, 15 ml of cells (3×10^6^/ml) were transfected with 15 μg of plasmid complexed with 30 μl Expifectamine293 transfection reagent. After 3 days the cells were harvested by centrifugation at 1000xg and after washing the pellets with PBS the cells were lysed in 1 ml cell lysis buffer/100 mg wet weight of cell pellet. After gentle shaking for 10 minutes the lysates were cleared of cell debris by centrifugation for 15 min at 14,000xg at 4º. Aliquots were stored at -20º or applied to either anti-FG or anti-HA agarose beads as described below or in the figure legends.

### Protein purification on affinity agarose beads

Beads were washed sequentially with 10 volumes TBS pH7.4 (to remove glycerol storage buffer), 10 volumes 0.1M glycine-HCL (to clear the antibody binding sites), and 10 volumes TBS, pH 7.4 to remove glycine-HCl and restore the pH to 7.4. Equal volumes of beads and cleared lysate were mixed by rotating 2-4 h or overnight at 4º. The slurry was added to a gravity flow disposable column, which was washed with 10 bed volumes of TBS-10 μM dT_52_, collecting 1 ml fractions, and then eluted with TBS-10 μM dT_52_ containing either 100 ug/ml 3XFG peptide or 500 μg/ml HA peptide in TBS-10 μM dT_52_ collecting 1 ml fractions. For the experiments shown in Figs. 3 and 4, we added the lysate bead slurry to Bio-Rad Micro Bio-Spin columns and washed the beads and eluted the bound proteins using a batch procedure: The agarose beads were resuspended in 1 ml wash buffer and recovered by centrifugation. The beads were then repeatedly washed with 1 ml elution buffer, collected by centrifugation saving the washes for further analysis. The bound proteins were eluted by suspending the beads in 1 ml of either of the above peptide-containing elution buffers. The beads were regenerated with 0.1 M glycine-HCl, pH 3.5.

### Western blots

Samples (25-50/μg) were added to 0.25 volume of NuPAGE 4X LDS Sample Buffer containing 20 mM DTT, heated for 10 min at 70º, and loaded onto a 1.0 mm NuPAGE 10% Bis-Tris polyacrylamide gel. After electrophoresis under denaturing conditions in NuPAGE™ MOPS 0.1% SDS Running Buffer (200 V, ∼40 min), the proteins were transferred to PVDF membranes using an iBlot. This instrument and all the reagents are Invitrogen™ products. The PVDF membranes were blocked with Thermo-Fisher SuperBlock T20 in 1x TBS and challenged with Abcam rabbit anti-beta tubulin polyclonal antibody and either Sigma-Aldrich mouse monoclonal anti-FG or Sigma-Aldrich mouse monoclonal anti-HA, overnight on a rocker at 4º. The PVDF membranes were washed 3 times (10 minutes/wash on a rocker) with 1xTBS/0.05% Tween-20 and then challenged with Sigma-Aldrich HRP-anti-mouse IgG for 1.5 hours on a rocker at room temperature. The membranes were washed as described above, and then 4 times with ∼25 ml 1x TBS to remove the Tween-20. Following the rinses, the membranes were incubated with Thermo-Fisher SuperSignal Pico West chemiluminescent substrate; the blots were exposed to film and developed.

### Mixed lysate experiment

We prepared cleared lysates as described above from cultures of HEK293F cells that had been separately transfected with pcDNA3.1-111-3XFG DNA or pcDNA3.1-151-HA DNA. Forty μl samples from each cleared lysate were combined and mixed with 40 μl of wash buffer (1x TBS-10 μM dT_52_). Also, forty μl of cleared lysate from either ORF1-111-3XFG or ORF1-151-HA transfected cells were diluted with 80 μl wash buffer. The three solutions were incubated at 25° C for thirty minutes, with occasional mixing. Thereupon the samples were transferred to 1.5 ml tubes containing 100 μl of Anti-FG M2 agarose beads (Millipore-Sigma) and incubated at 25° C for 2 hours, with constant mixing in an Eppendorf Thermomixer R, after which the tubes were centrifuged at 1000xg for 5 minutes. The binding step supernatants were saved and the agarose beads were collected by centrifugation, resuspended in wash buffer and centrifuged for 5 minutes at 1000xg. The wash was repeated four times saving the wash supernatants. Protein bound by the anti-Flag M2 agarose beads was eluted by incubation in 100 ul elution buffer (1x-TBS-10 μM dT_52_, containing 100 ug/ml 3XFG peptide) for 30 minutes on a Thermomixer shaker. After incubation, the bead slurry was transferred to a Micro Bio-spin columns and the eluate was collected by a 1 min centrifugation at 1000xg. This elution step was carried out three times. The binding step supernatants, wash supernatants and eluates were evaluated by western blot as described above.

## Acknowledgement

The authors thank Nicholas Guydosh (LBG/NIDDK/NIH) for his informative comments and suggestions.

## Notes

### Competing Interest Statement

The authors have declared no competing interest.

