## Supplemental Information for "CO-EXPRESSION OF DISTINCT COILED COILS CAN LEAD TO THEIR ENTANGLEMENT"

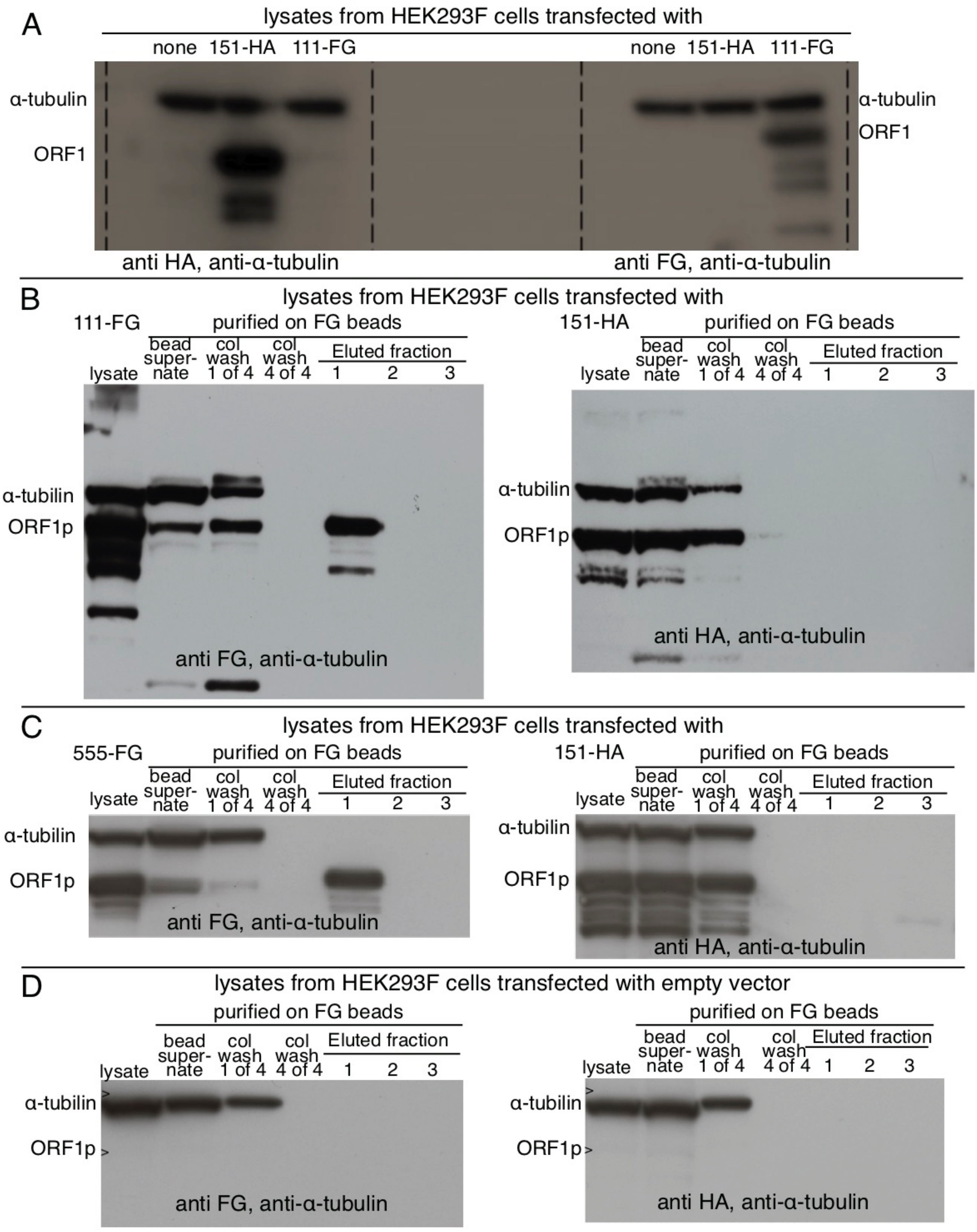

Supplemental Figure 1.

Control experiments that support the conclusions drawn from Figures 2-4 of the main paper, : Panel A show robust expression of both tagged proteins; Panels B and C show that only FG-tagged proteins bind to the anti FG-beads, and panel D shows that lysates from cells transfected with the empty pcDNA.1(+) expression vector lack any proteins that cross react with the anti-FG or anti-FA antibodies used to develop the Western blots.

>555-1xFG
ggatccgcaATGGGGAAAAAACAGAGCAGAAAAGCTGAAAATTCTAAAAATCAGAGCGCCTCTCCTCCTCCAAAGGA
ACGCAGCTCCTCACCAGCAACGGAACAAAGCTGGATGGAGAATGACTTTGACGAGTTGAGAGAAGAAGGCTTCAGAC
GATCAAACTTCTCCGAGCTAAAGGAGGAAGTTCGAACCCATCGCAAAGAAGCTAAAAACCTTGAAAAAAGATTAGAC
GAATGGCTAACTAGAATAACCAGTGTAGAGAAGTCCTTAAATGACCTGATGGAGCTGAAAACCATGGCACGAGAACT
ACGTGACGAATGCACAAGCTTCAGTAGCCGATTCGATCAACTGGAAGAAAGGGTATCAGTGATTGAAGATCAAATGA
ATGAAATGAAGCGAGAAGAGAAGTTTAGAGAAAAAAGAGTAAAAAGAAATGAACAAAGCCTCCAAGAAATATGGGAC
TATGTGAAAAGACCAAATCTACGTCTGATTGGTGTACCTGAAAGTGACGGGGAGAATGGAACCAAGTTGGAAAACAC
TCTGCAGGATATTATCCAGGAGAACTTCCCCAACCTAGCAAGGCAGGCCAACATTCAAATTCAGGAAATACAGAGAA
CGCCACAAAGATACTCCTCGAGAAGAGCAACTCCAAGACACATAATTGTCAGATTCACCAAAGTTGAAATGAAGGAA
AAAATGTTAAGGGCAGCCAGAGAGAAAGGTCGGGTTACCCACAAAGGGAAGCCCATCAGACTAACAGCGGATCTCTC
GGCAGAAACTCTACAAGCCAGAAGAGAGTGGGGGCCAATATTCAACATTCTTAAAGAAAAGAATTTTCAACCCAGAA
TTTCATATCCAGCCAAACTAAGCTTCATAAGTGAAGGAGAAATAAAATCCTTTACAGACAAGCAAATGCTGAGAGAT
TTTGTCACCACCAGGCCTGCCCTACAAGAGCTCCTGAAGGAAGCACTAAACATGGAAAGGAACAACCGGTACCAGCC
ACTGCAAAAACATGCCAAATTGgattataaggacgacgacgacaagtaggaattc
>111-3XFG
ggatccgcaATGGGGAAAAAACAGAACAGAAAAACTGGAAACTCTAAAACGCAGAGCGCCTCTCCTCCTCCAAAGGA
ACGCAGTTCCTCACCAGCAACaGAACAAAGCTGGATGGAGAATGATTTTGACGAGCTGAGAGAAGAAGGCTTCAGAC
GATCAAATTACTCTGAGCTACGGGAGGACATTCAAACCAAAGGCAAAGAAGTTGAAAACTTTGAAAAAAATTTAGAA
GAATGTATAACTAGAATAACCAATACAGAGAAGTGCTTAAAGGAGCTGATGGAGCTGAAAACCAAGGCTCGAGAACT
ACGTGAAGAATGCAGAAGCCTCAGGAGCCGATGCGATCAACTGGAAGAAAGGGTATCAGCAATGGAAGATGAAATGA
ATGAAATGAAGCGAGAAGGGAAGTTTAGAGAAAAAAGAATAAAAAGAAATGAGCAAAGCCTCCAAGAAATATGGGAC
TATGTGAAAAGACCAAATCTACGTCTGATTGGTGTACCTGAAAGTGATGTGGAGAATGGAACCAAGTTGGAAAACAC
TCTGCAGGATATTATCCAGGAGAACTTCCCCAATCTAGCAAGGCAGGCCAACGTTCAGATTCAGGAAATACAGAGAA
CGCCACAAAGATACTCCTCGAGAAGAGCAACTCCAAGACACATAATTGTCAGATTCACCAAAGTTGAAATGAAGGAA
AAAATGTTAAGGGCAGCCAGAGAGAAAGGTCGGGTTACCCTCAAAGGAAAGCCCATCAGACTAACAGTGGATCTCTC
GGCAGAAACCCTACAAGCCAGAAGAGAGTGGGGGCCAATATTCAACATTCTTAAAGAAAAGAATTTTCAACCCAGAA
TTTCATATCCAGCCAAACTAAGCTTCATAAGTGAAGGAGAAATAAAATACTTTATAGACAAGCAAATGTTGAGAGAT
TTTGTCACCACCAGGCCTGCCCTAAAAGAGCTCCTGAAGGAAGCgCTAAACATGGAAAGGAACAaccggtACCAGCC
GCTGCAAAATCATGCCAAAATGgactacaaagaccatgacggtgattataaagatcatgacatcgattacaaggatg
acgatgacaagTAGgaattc
>151-HA
ggatccgcAATGGGGAAAAAACAGAACAGAAAAACTGGAAACTCTAAAACGCAGAGCGCCTCTCCTCCTCCAAAGGA
ACGCAGTTCCTCACCAGCAACAGAACAAAGCTGGATGGAGAATGATTTTGACGAGCTGAGAGAAGAAGGCTTCAGAC
GATCAAATTACTCTGAGCTACGGGAGGACATTCAAACCAAAGGCAAAGAAGTTGAAAACTTTGAAAAAAATTTAGAA
GAATGTATAACTAGAATAACCAATACAGAGAAGTGCTTAAAGGAGCTGATGGAGCTGAAAACCAAGGCTCGAGAACT

ACGTGAAGAATGCACAAGCTTCAGCAGCCGATTCGATCAACTGGAAGAAAGGGTATCAGTTATCGAAGATCAAATGA
ATGAAATGAAGCGAGAAGAGAAGTTTAGAGAAAAAAGAGTAAAAAGAAATGAGCAAAGCCTCCAAGAAATATGGGAC
TATGTGAAAAGACCAAATCTACGTCTGATTGGTGTACCTGAAAGTGATGTGGAGAATGGAACCAAGTTGGAAAACAC
TCTGCAGGATATTATCCAGGAGAACTTCCCCAATCTAGCAAGGCAGGCCAACGTTCAGATTCAGGAAATACAGAGAA
CGCCACAAAGATACTCCTCGAGAAGAGCAACTCCAAGACACATAATTGTCAGATTCACCAAAGTTGAAATGAAGGAA
AAAATGTTAAGGGCAGCCAGAGAGAAAGGTCGGGTTACCCTCAAAGGAAAGCCCATCAGACTAACAGTGGATCTCTC
GGCAGAAACCCTACAAGCCAGAAGAGAGTGGGGGCCAATATTCAACATTCTTAAAGAAAAGAATTTTCAACCCAGAA
TTTCATATCCAGCCAAACTAAGCTTCATAAGTGAAGGAGAAATAAAATACTTTATAGACAAGCAAATGTTGAGAGAT
TTTGTCACCACCAGGCCTGCCCTAAAAGAGCTCCTGAAGGAAGCGCTAAACATGGAAAGGAACAACCGGTACCAGCC
GCTGCAAAATCATGCCAAAATGTACCCATACGATGTTCCAGATTACGCTgTAGgaattc

Supplemental Figure 2 - ORF1. sequences flanked by BamHI and EcoRI recognition sites
